# BACTERIAL AND FUNGAL CONTAMINATION OF STAIRCASE BANISTERS AT THE COLLEGE OF SCIENCE, KWAME NKRUMAH UNIVERSITY OF SCIENCE AND TECHNOLOGY, GHANA

**DOI:** 10.64898/2026.08.07.743594

**Authors:** Evans Akwaboah, Bernice Awotwe-Mensah, Francisca Obeng-Mensah, Richard Frimpong Koranteng, Anokyewaa Abigail Appau, Emmanuel Ndezure, Linda Aurelia Ofori

## Abstract

Staircase banisters are frequently touched surfaces that may receive microorganisms from hands, dust, air and other environmental sources, but their microbial status in Ghanaian university buildings has received limited attention. This cross-sectional environmental microbiology study assessed bacterial and fungal contamination of staircase banisters at the Kwame Nkrumah University of Science and Technology, Kumasi. Six banisters from the Aboagye Menyah Building Complex, Chemistry Block and Biology Block were purposively selected to include high-traffic locations and both wooden and metal surfaces. Upper and lower sections were sampled over three consecutive Monday afternoons after classes, giving 12 surface samples. Approximately 150 cm² of each section was swabbed with sterile buffered peptone water, cultured on standard bacteriological and mycological media, and analysed using phenotypic and morphological methods. Bacterial loads were compared by independent samples t-test. Thirty-one bacterial isolates were recovered. The study found Gram-positive bacteria which accounted for 74% of isolates and Gram-negative bacteria for 26%. *Staphylococcus spp.*, *Streptococcus spp.*, *Enterobacter*iaceae, *Bacillus spp.* and *Corynebacterium-+ spp.* were the main presumptive bacterial groups. Metal banisters had higher mean bacterial loads than wooden banisters (4.38 ± 0.86 versus 1.24 ± 1.44 log10 CFU/mL; p = 0.014), whereas upper and lower sections did not differ significantly (p = 0.539). Fungal growth was detected in all samples, with *Aspergillus fumigatus*, *Colletotrichum spp.* and *Aspergillus niger* being frequent presumptive fungi. The findings support the routine inclusion of staircase banisters in cleaning and disinfection programmes for academic buildings.

## INTRODUCTION

Microorganisms are pervasive across natural and built environments, and indoor microbial communities are continuously shaped by human occupancy, outdoor air, dust, water, building materials, ventilation, and routine surface contact (Gilbert & Hartmann, 2024; Zampolli *et al*., 2024). Indoor surfaces are important because people spend much of their time inside buildings and repeatedly touch objects such as door handles, worktops, mobile devices, railings and staircase banisters (Klepeis *et al*., 2001; Stephens *et al*., 2019). When contaminated with viable microorganisms, these inanimate surfaces may act as fomites and support microbial exchange between human hands, environmental surfaces and other objects. High-touch surfaces are therefore important interfaces between routine human activity and environmental microbiology (Cobrado *et al*., 2017; Appiah *et al*., 2025).

The persistence of microorganisms on surfaces depends on organism type, initial microbial load, surface material, temperature, humidity, organic matter and cleaning practices. Bacteria, fungi, protozoa and viruses may retain replication capacity on inanimate surfaces for variable periods, although laboratory persistence data often represent worst-case conditions and should be interpreted carefully for real-world settings (Kramer *et al*., 2024). Surface-associated microorganisms may also occur in biofilms, which can increase tolerance to desiccation and disinfectants (Donlan, 2002; Mirghani *et al*., 2022). This persistence matters because contaminated surfaces may contribute to indirect microbial transfer through hand contact, especially where many users touch the same surface within short periods.

Staircase banisters provide support, balance and stability during movement between floors (Ishihara *et al*., 2002). In academic buildings, the same banister may be touched repeatedly by students, staff, visitors and cleaners. Transfer can occur in both directions: users may deposit skin-associated, respiratory or environmental organisms onto a banister, and they may also pick up organisms already present on the surface. Experimental work with fluorescent tracers has shown that contamination acquired from public surfaces can be transferred to hands, personal belongings and domestic environments (Reynolds *et al*., 2005). Banisters may also receive dust, bacterial aerosols and fungal spores from indoor and outdoor air (Fröhlich-Nowoisky *et al*., 2016; Gołofit-Szymczak and Górny, 2010).

Direct research on staircase handrails and banisters remains limited compared with studies on other high-touch surfaces. In Egypt, bacterial contamination was reported on primary school surfaces, including banisters, with *Staphylococcus spp.* among the common isolates (El-Kased and Gamaleldin, 2020). In Zambia, Mulongo *et al*. (2021) recovered *Staphylococcus aureus*, coagulase-negative staphylococci, Gram-positive bacilli, streptococci, *Klebsiella spp.* and *Enterobacter spp.* from elevators and staircase handrails in a teaching hospital. Elevator buttons in university buildings have also been reported to carry bacterial and fungal contamination (Mohammadi *et al*., 2016). More recent campus microbiome evidence shows that bacterial communities on university surfaces can be traced substantially to building use and human skin sources (Ye *et al*., 2024), while high-touch surfaces in university and urban settings may carry organisms with potential pathogenic and antimicrobial resistance relevance (Liu *et al*., 2026).

Contamination patterns may differ between healthcare and university environments. Healthcare facilities usually have higher proportions of vulnerable users and more formal infection prevention protocols, while universities combine high population movement, variable hand hygiene, variable cleaning frequency and frequent contact with lecture rooms, laboratories, offices and stairways. Therefore, findings from hospitals cannot be transferred directly to academic settings, especially in Ghana where evidence on staircase banisters and related university high-touch surfaces is still limited.

Surface material may also influence microbial retention and survival. Porosity, moisture retention, surface roughness, residual organic matter and biofilm formation can affect how microorganisms attach to wood or metal. Some wood surfaces may absorb moisture or contain natural antimicrobial compounds, while non-porous metal surfaces may retain residues and moisture films that support surface attachment under certain conditions (Milling *et al*., 2005; Munir *et al*., 2019; Chen *et al*., 2020). Comparing wooden and metal banisters is therefore relevant for sanitation planning.

This study aimed to assess bacterial and fungal contamination of selected staircase banisters at the College of Science, Kwame Nkrumah University of Science and Technology, Ghana. The specific objectives were to isolate and presumptively identify bacterial and fungal contaminants from staircase banisters, compare bacterial loads between wooden and metal banisters, and compare bacterial loads between upper and lower sections of the banisters.

## MATERIALS AND METHODS

### Study design and setting

This was a cross-sectional environmental microbiology study conducted from January to August 2024 at the College of Science, Kwame Nkrumah University of Science and Technology, Kumasi, Ghana. Sampling was performed in the Aboagye Menyah Building Complex, Chemistry Block and Biology Block. Laboratory processing was undertaken in the Microbiology Laboratory of the Department of Theoretical and Applied Biology.

Surface sampling was carried out over three consecutive Monday afternoons after classes. This period was selected because student and staff movement through lecture rooms, laboratories, offices and assignment submission areas was expected to be high, increasing the relevance of the sampled surfaces as frequently touched fomites.

### Sampling sites and sample size

Six staircase banisters were purposively selected to represent frequently used staircase routes within the College of Science and to include both wooden and metal surfaces. Selection considered expected human traffic, building relevance, accessibility for sampling and material type. One relatively isolated metal banister was also included among the sampled sites; however, the study was not designed or powered to compare isolated and high-traffic banisters as separate exposure groups.

Three banisters were sampled from the Aboagye Menyah Building Complex, one from the Chemistry Block and two from the Biology Block. Each banister was divided into upper and lower sampling sections, giving 12 surface samples in total. The sample coding system is shown in Table I.

**Table I.**
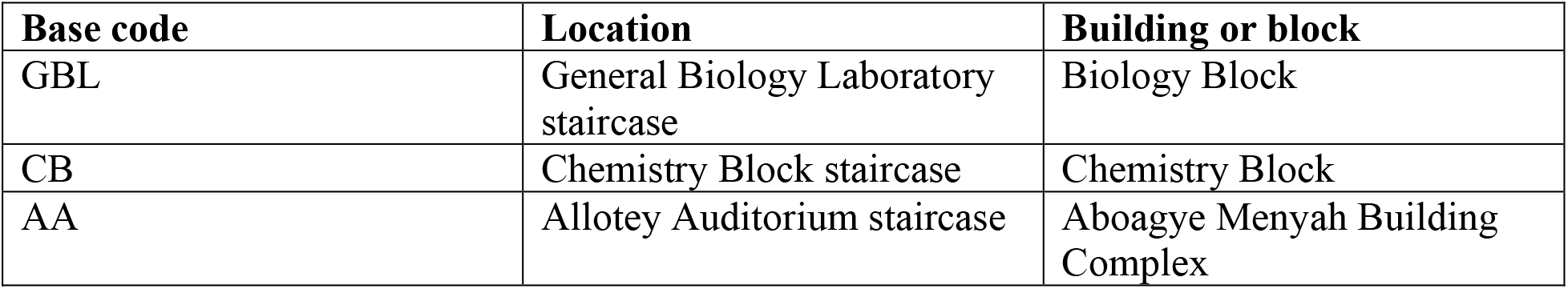
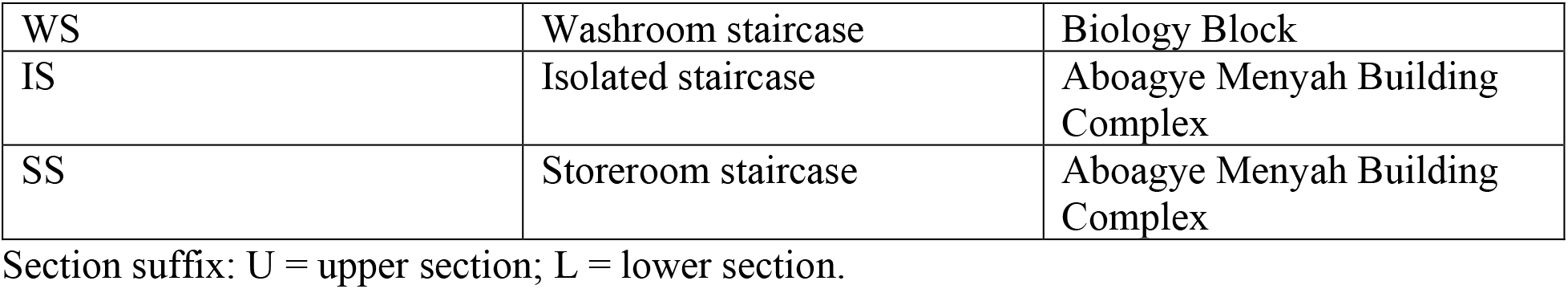
Sampling codes and locations of selected staircase banisters.

### Sample collection

Sterile cotton swabs, gloves, sample bags and laboratory materials were prepared before sampling. At each site, a sterile swab was moistened with sterile buffered peptone water and rubbed across an approximately 30 cm by 5 cm area of the banister surface, equivalent to about 150 cm². The swab was moved both horizontally and vertically to improve surface contact. Each swab was returned to a labelled sterile bag, sealed, placed in an ice chest and transported to the laboratory for processing.

### Sample preparation, microbial culture and isolation

In the laboratory, 9 mL of sterile buffered peptone water was added to each bag containing a swab. The sample was mixed in a pulsifier for 30 seconds to produce the stock suspension. Tenfold serial dilutions were prepared to 10−3. Cultures were prepared in triplicate where applicable, and the average count was used to calculate the microbial count for each banister section.

Plate Count Agar was used to estimate total viable bacterial counts because it supports the growth of a broad range of heterotrophic bacteria. Mannitol Salt Agar was used for the selective and differential recovery of staphylococci, MacConkey Agar for Gram-negative enteric bacteria, Nutrient Agar for purification of bacterial isolates and Potato Dextrose Agar for fungal isolation. One milliliter of the selected dilution was inoculated into labelled Petri dishes. Bacterial plates were incubated at 37 °C for 24 hours, while fungal plates were monitored until visible colonies developed after approximately 48 hours of incubation.

### Bacterial enumeration and identification

Countable bacterial colonies were enumerated with a colony counter and expressed as colony-forming units per millilitre. The count was calculated from the average colony count, dilution factor and plated volume. Bacterial loads were summarized on a log10 CFU/mL scale. Distinct colonies were subcultured on Nutrient Agar to obtain pure cultures. Presumptive identification was based on colony appearance, Gram staining, cell morphology and biochemical reactions. The biochemical tests included catalase, citrate utilization, indole production and glucose and lactose fermentation. Because the study used phenotypic methods only, bacterial assignments are reported as presumptive genera or groups.

### Fungal enumeration and identification

Fungal colonies were counted and expressed as CFU/mL based on the average colony count from replicated cultures and serial dilutions. Fungal cultures were examined by macroscopic colony morphology, including colour, texture, surface appearance, margin and growth form. A hand lens and stereomicroscope were used where necessary. Microscopic examination was performed using lactophenol cotton blue wet mounts, a standard staining method for visualizing fungal hyphae and reproductive structures (Leck, 1999). The plates were also reviewed with mycological assistance. Fungal genus and species assignments are therefore reported as presumptive because molecular confirmation was not performed.

### Statistical analysis

Data were entered and summarized in Microsoft Excel LTSC 2021 and analysed in IBM SPSS version 27. Means, standard deviations, ranges, frequencies and percentages were calculated. Bacterial loads were compared between wooden and metal banisters and between upper and lower banister sections using independent samples t-tests. A p value below 0.05 was considered statistically significant. The small number of sampled banisters was considered when interpreting statistical comparisons.

## RESULTS

### Bacterial isolates and distribution

Thirty-one bacterial isolates were recovered from the 12 banister samples. Based on broad Gram reaction, Gram-positive bacteria accounted for 74% of the isolates and Gram-negative bacteria for 26%. For the organism groups with complete phenotypic assignment, the dominant presumptive bacterial group was *Staphylococcus spp.* (10/31; 32.3%), followed by *Streptococcus spp.* (7/31; 22.6%), *Enterobacter*iaceae (6/31; 19.4%), *Bacillus spp.* (5/31; 16.1%) and *Corynebacterium spp.* (2/31; 6.5%). One isolate was not assigned beyond the available broad classification record (Figure 1).

**Figure 1.**
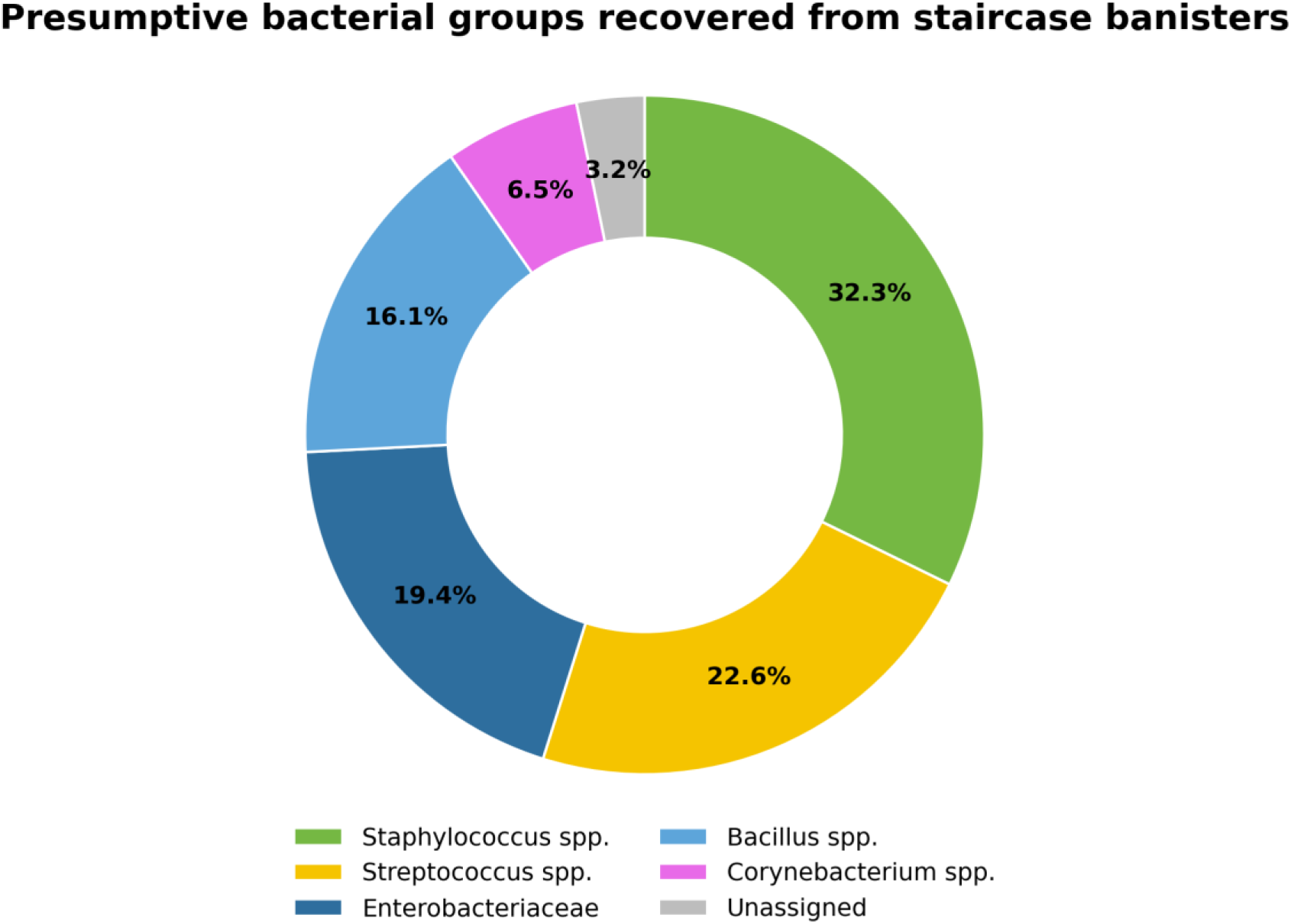
Distribution of presumptive bacterial groups recovered from staircase banisters. Percentages are based on the 31 bacterial isolates recovered; one isolate was not assigned beyond the available broad classification record.

### Bacterial load across banisters

Bacterial load varied among the sampled banister sections (Figure 2). No countable bacterial growth was recorded for GBLU and CBL under the culture conditions used. The highest sample-level loads were recorded at WSL and SSU. The mean load for upper sections was 3.17 ± 1.89 log10 CFU/mL, while the mean load for lower sections was 3.50 ± 1.94 log10 CFU/mL. This difference was not statistically significant using an independent samples t-test (p = 0.539). Metal banisters had a higher mean bacterial load than wooden banisters (4.38 ± 0.86 versus 1.24 ± 1.44 log10 CFU/mL), and this difference was statistically significant using an independent samples t-test (p = 0.014) (Table II).

**Figure 2.**
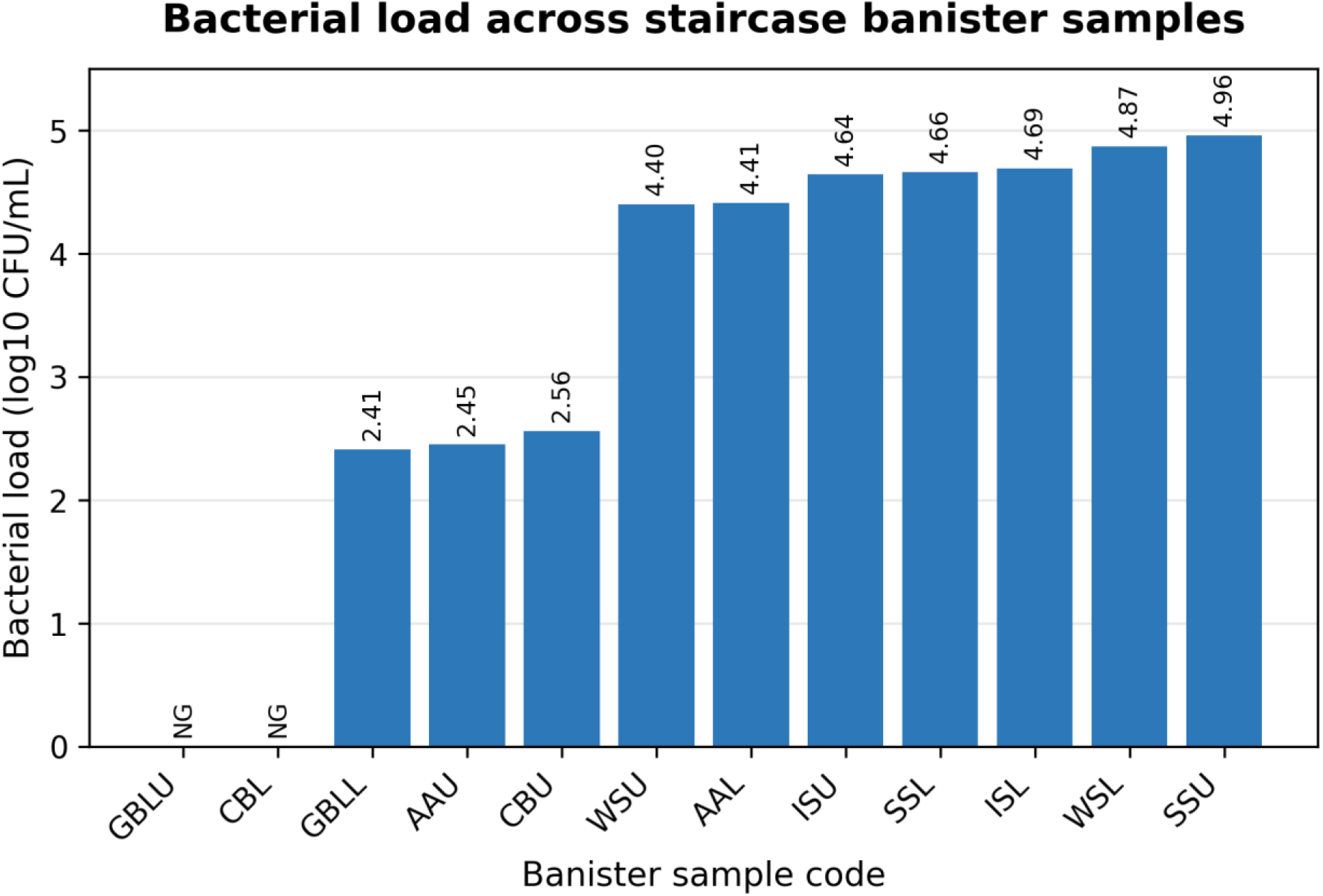
Bacterial load across the 12 staircase banister samples. Values are shown as log10 CFU/mL. NG indicates that no countable growth was recorded under the culture conditions used. Sample codes are defined in Table I.

**Table II.** Bacterial load according to banister position and surface material.

| Comparison | Group | Mean log<sub>10</sub> CFU/mL (SD) | p value |
| --- | --- | --- | --- |
| Position | Upper section | 3.17 (±1.89) | 0.539 |
|  | Lower section | 3.50 (±1.94) |  |
| Material | Wooden banister | 1.24 (±1.44) | 0.014 |
|  | Metal banister | 4.38 (±0.86) |  |

p values were obtained using independent samples t-tests. The p values are reported from the original thesis analysis dataset used for the group comparisons.

### Fungal load and distribution

Fungal growth was detected in all 12 sampled sections. Fungal CFU/mL values were calculated from averaged colony counts after serial dilution. AAU and CBU had the highest fungal counts, with 13 CFU/mL each, followed by ISL with 11 CFU/mL (Figure 3). In total, 66 fungal colonies were classified. The predominant presumptive fungus was *Aspergillus fumigatus* (30/66; 45.5%). Other presumptive fungi included *Colletotrichum spp.* (14/66; 21.2%), *Aspergillus niger* (10/66; 15.2%), *Neurospora spp.* (6/66; 9.1%), *Aspergillus flavus* (3/66; 4.5%) and *Rhizopus spp.* (3/66; 4.5%) (Figure 4).

**Figure 3.**
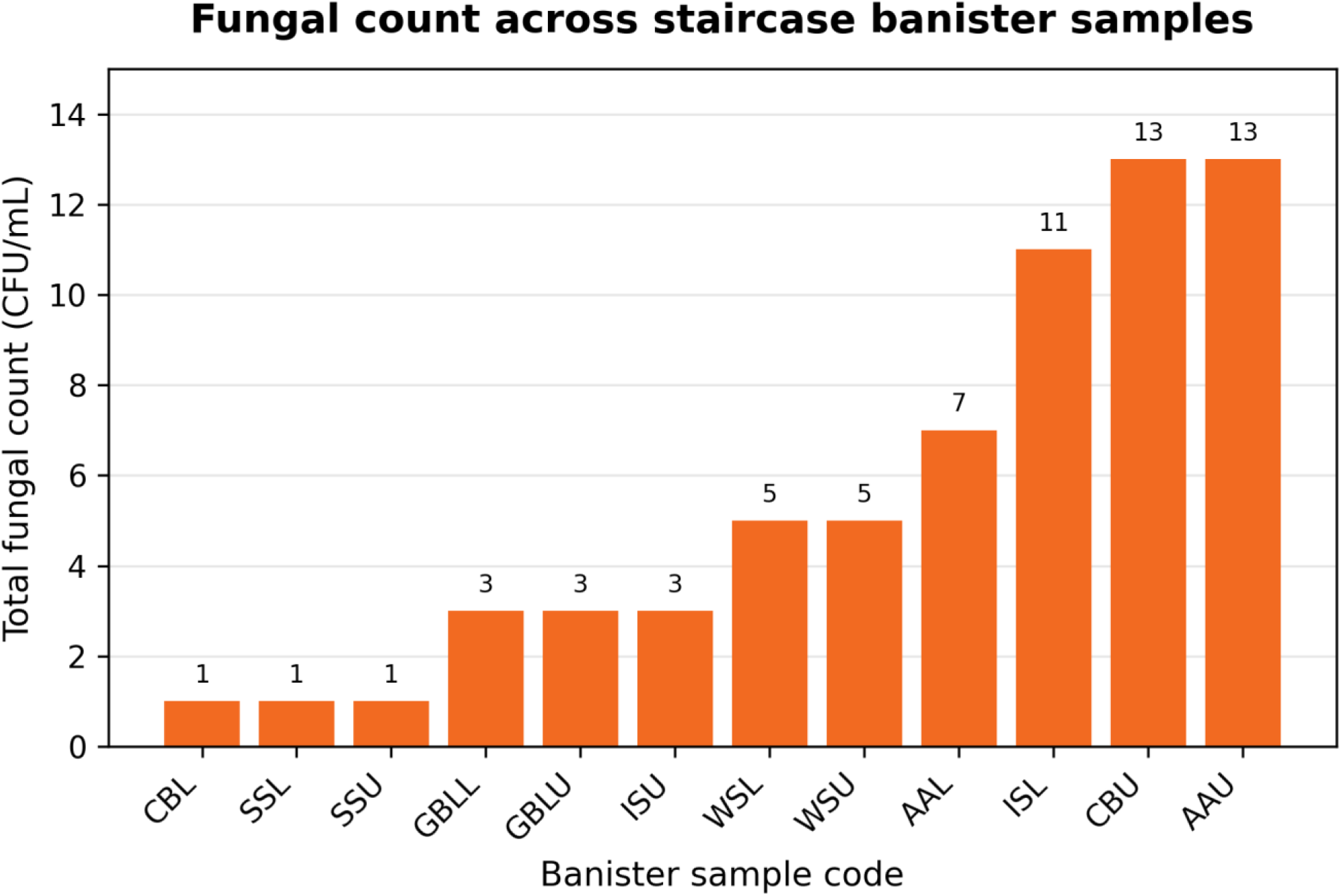
Total fungal counts across the 12 staircase banister samples. Values are expressed as CFU/mL after averaging colony counts from replicated cultures and applying the dilution factor. Sample codes are defined in Table I.

**Figure 4.**
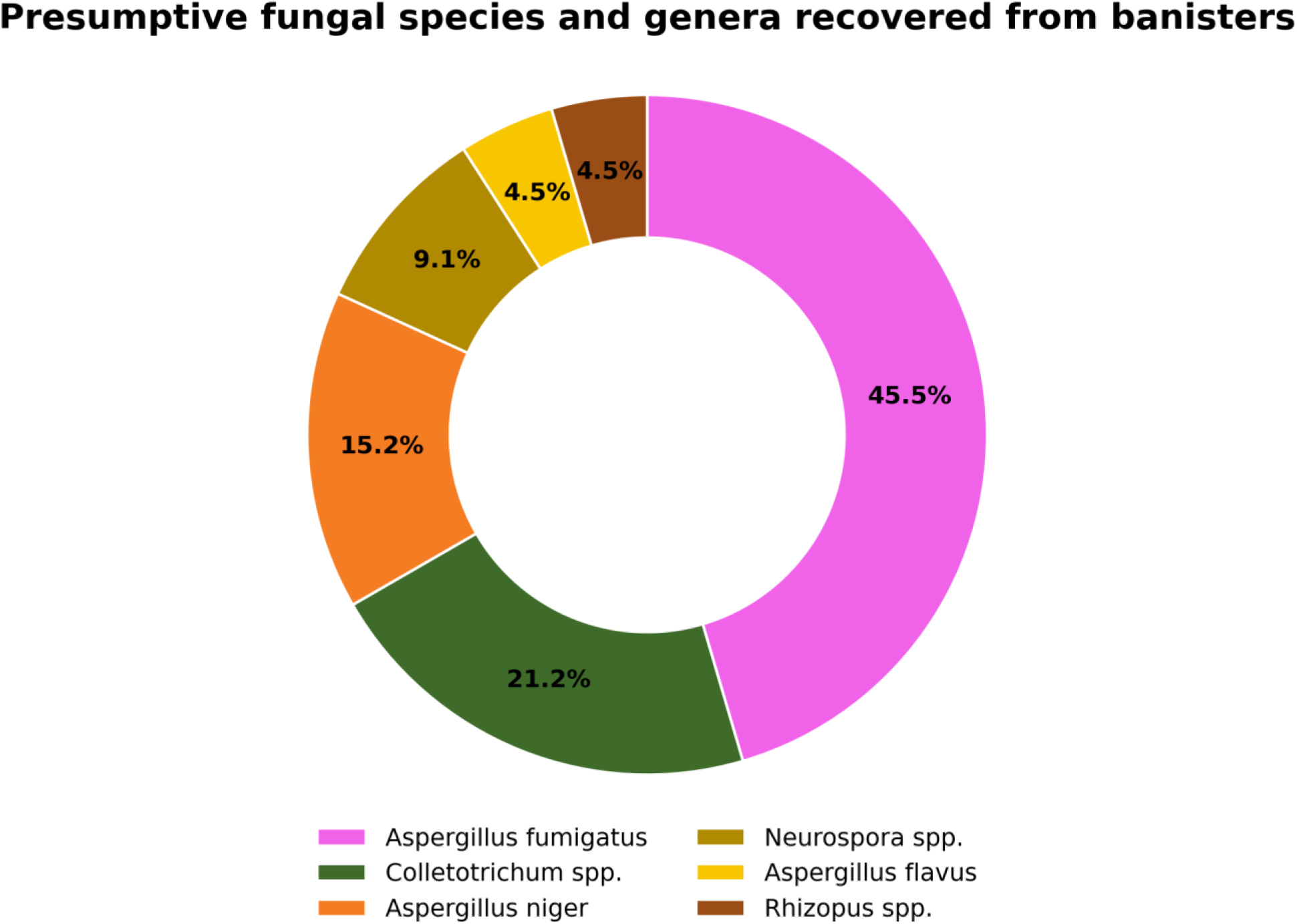
Distribution of presumptively identified fungal species and genera recovered from the sampled banisters (n = 66 fungal colonies).

## DISCUSSION

This study found bacterial and fungal contamination on staircase banisters in three buildings within the College of Science, Kwame Nkrumah University of Science and Technology. The recovered organisms included skin-associated bacteria, respiratory and environmental bacteria, enteric bacterial groups and airborne or environmental fungi. Bacterial loads differed by surface material, with metal banisters showing a higher observed mean bacterial load than wooden banisters. In contrast, upper and lower sections did not differ significantly. These findings support the view that staircase banisters are overlooked high-touch fomites in academic buildings (Silas *et al*., 2026).

The recovery of microorganisms from the sampled banisters is consistent with studies of handrails, elevator buttons, door handles and other frequently touched surfaces. Mulongo *et al*. (2021) recovered diverse bacteria from staircase handrails and elevators in a teaching hospital, while Mohammadi *et al*. (2016) reported bacterial and fungal contamination on elevator buttons in university buildings. A recent systematic review of door handles also confirms that high-touch public and healthcare surfaces may carry bacterial, fungal and viral contaminants (Appiah *et al*., 2025). The present study did not measure organism transfer to users or clinical outcomes. Detection on a banister therefore does not prove infection transmission, but it identifies a point at which microorganisms may be exchanged between hands and the built environment.

The higher observed bacterial load on metal banisters should be interpreted cautiously because only a small number of wooden and metal banisters were sampled and potential confounders such as human traffic, cleaning frequency, surface age, surface roughness and nearby activities were not measured. Nevertheless, surface material is biologically plausible as a factor affecting microbial retention and survival. Wood can absorb moisture into its structure and may contain natural antimicrobial compounds, while non-porous metal surfaces can retain organic residues and moisture films that support attachment under some conditions (Milling *et al*., 2005; Munir *et al*., 2019; Chen *et al*., 2020). Biofilm formation on hard surfaces may further support persistence and resistance to routine cleaning (Donlan, 2002; Mirghani *et al*., 2022).

The absence of a significant difference between upper and lower banister sections suggests that contamination was not restricted to one vertical section. Users may touch different points depending on direction of movement, height, balance, crowding and personal habits. Dust and bioaerosols can also settle along the length of a banister. The wide variability within both positions indicates that location-specific factors may have had a stronger influence than section position alone. The small sample size may also have limited statistical power to detect a true difference between upper and lower sections.

*Staphylococcus spp.* formed the largest presumptive bacterial group. *Staphylococci* are common members of skin microbiota and may be transferred readily by hand contact. This agrees with Mulongo *et al*. (2021), who reported *Staphylococcus aureus* and coagulase-negative *staphylococci* among organisms recovered from staircase handrails and elevators. *Streptococcus* spp. were also recorded and may reflect contact with oral or respiratory secretions, contaminated hands or environmental droplets (Santacroce *et al*., 2020). *Bacillus* spp. are environmentally widespread and can persist through spore formation, while *Corynebacterium* spp. are commonly associated with human skin, although some members may act as opportunistic pathogens (Bernard, 2005).

Enterobacteriaceae represented a notable proportion of the presumptive bacterial isolates. This finding should not be interpreted as definitive evidence of faecal contamination because species-level identification and source tracking were not performed. Possible explanations include inadequate hand hygiene, indirect transfer from objects, dust deposition, environmental contamination or human-mediated transfer. The phenotypic methods used in this study were not sufficient to resolve all isolates reliably to species level. The public health importance of individual isolates therefore cannot be determined without confirmatory identification and antimicrobial susceptibility testing.

Fungal contamination was also widespread. *Aspergillus fumigatus* was the most frequent presumptive species, while *Aspergillus niger*, *Aspergillus flavus*, *Neurospora spp.*, *Rhizopus spp.* and *Colletotrichum spp.* were also recorded. Indoor and outdoor fungal profiles are influenced by ventilation, season, moisture, dust and surrounding vegetation (Gołofit-Szymczak and Górny, 2010; Sautour *et al*., 2009). *Aspergillus fumigatus* is an important environmental mould and can cause disease in susceptible people, particularly those with impaired immunity (Brakhage and Langfelder, 2002). *Colletotrichum spp.* are strongly associated with plants and may plausibly reflect deposition from nearby vegetation or airborne plant material (Talhinhas and Baroncelli, 2021). Because fungal identification relied on macroscopic and microscopic morphology, molecular confirmation would be required for definitive species identification.

The findings have practical sanitation implications for academic buildings. Staircase banisters should be recognized in written cleaning schedules as high-touch surfaces, alongside door handles, desks, railings and laboratory contact points. Cleaning should remove visible soil and organic material before disinfectant application at appropriate concentration and contact time. However, the present study did not test cleaning interventions, so recommendations are based on contamination evidence rather than measured disinfection effectiveness. Hand-hygiene awareness can provide an additional barrier by reducing microbial deposition and transfer.

This study has several strengths. It examined both bacterial and fungal contamination, sampled multiple College of Science buildings, compared two surface materials and evaluated upper and lower banister sections. It also provides baseline evidence from a Ghanaian university environment, where data on staircase banisters are scarce. Limitations include the small sample size, sampling within one institution, absence of repeated seasonal sampling, lack of measured traffic and cleaning history, limited statistical power and culture-based detection only. The study also relied on phenotypic and morphological identification without molecular confirmation or antimicrobial susceptibility testing. Future studies should use larger sample sizes, repeated sampling across seasons and time points, standardized surface templates, recorded cleaning history, molecular identification and hand-transfer experiments.

## CONCLUSIONS

The study uncovered a variety of bacterial and fungal contamination on staircase banisters at the College of Science, Kwame Nkrumah University of Science and Technology, Kumasi. While these numbers are relatively low, these findings support the routine inclusion of staircase banisters in cleaning and disinfection programmes for academic buildings and highlight the need for larger studies using confirmatory identification methods and more detailed exposure assessment

## ACKNOWLEDGEMENTS

The authors acknowledge Mr Acheampong and the laboratory technicians of the Department of Theoretical and Applied Biology for technical and logistical support during the undergraduate research project.

## ETHICS APPROVAL

The study involved environmental surface sampling only and did not involve human participants, animals or identifiable personal data. Formal ethics approval was therefore not applicable.

## DATA AVAILABILITY

The data supporting the findings are presented within the manuscript and the underlying undergraduate thesis. Additional study records may be requested from the corresponding author.

## AUTHOR CONTRIBUTIONS

EA contributed to study conceptualization, project leadership, methodology, sampling, laboratory investigation, data curation, formal analysis, original drafting, manuscript revision and corresponding author responsibilities. BAM, FOM and RFK contributed as undergraduate project students to sampling, laboratory investigations, data curation and manuscript review. AAA co-supervised the work and provided microbiological technique support. EN contributed to manuscript writing, review and preparation. LAO supervised the research, guided interpretation and reviewed the manuscript. All authors reviewed and approved the final manuscript.

